# Perirhinal and postrhinal cortices are necessary for retrieving latently-acquired stimulus associations

**DOI:** 10.1101/2024.05.26.595958

**Authors:** Xiangyuan Peng, Rebecca D. Burwell

## Abstract

The perirhinal and postrhinal cortices reside in the parahippocampal region of the medial temporal lobe. They receive and process sensory information from cortical and subcortical sources and support hippocampal functions via direct connectivity and indirectly via the entorhinal cortex. Previous studies using the sensory preconditioning paradigm have shown that the perirhinal and postrhinal cortices are necessary for associating cues during preconditioning and retrieving the associations during conditioning. However, the question of whether these regions are also required for stimulus association retrieval during re-exposure to the preconditioning cue had not been addressed. Using a chemogenetic approach, we temporarily suppressed the perirhinal or postrhinal cortex in adult male rats during the preconditioning cue test phase. Both suppression groups showed impaired sensory preconditioning compared to sham surgery controls, as indicated by significantly reduced preferential responding. Implications of our findings are discussed in relation to the importance of the PER and POR in context processing and episodic-like memory in animal models.

**Significance statement:** Although Pavlovian conditioning between two stimuli is widely used in neuroscience, associative learning in the real-world is often too complex to be modeled by first-order conditioning alone. Proper responding may require integrating multiple associations via common elements, i.e. higher-order conditioning, the neural basis of which is not well understood. Whereas existing research indicates the perirhinal and postrhinal cortices in the medial temporal lobe contribute to certain forms of sensory preconditioning (SPC, a type of higher-order conditioning), we suggest their contributions lie in the encoding and retrieving latently acquired associations, processes which are central to all forms of SPC. We argue this interpretation also captures perirhinal and postrhinal contributions to contextual and episodic memory, hence offering a unifying explanation regarding the contributions of these cortices across multiple paradigms.

## Introduction

The medial temporal lobe (MTL) plays pivotal roles in a range of memory functions underpinning human and animal behavior. In the rodent MTL, the perirhinal cortex (PER) and postrhinal cortex (POR, homologous to the primate parahippocampal cortex) are critically involved in various forms of learning and memory (Burwell et al., 1995; Estela-Pro & Burwell, 2021; Peng & Burwell, 2021). Both regions receive polymodal and unimodal associational input from other cortical and subcortical structures (Burwell & Amaral, 1998a; Furtak et al., 2007; Tomas Pereira et al., 2016). Information from the PER and POR reaches hippocampal CA1 and the subiculum through direct projections and indirect connections via the entorhinal cortex (Agster & Burwell, 2013). The PER specializes in processing item features (i.e. “what”), whereas the POR mainly processes spatial properties of items and the environment ( i.e. “where”). Through robust, reciprocal connections, the PER and POR work together to represent the local environment, including the spatial layout of the items, objects, and patterns contained in the spatial context (Burwell & Amaral, 1998b; Estela, 2020; Furtak et al., 2012; Heimer-McGinn et al., 2017). The information is used by the hippocampus to support episodic, recognition, and other forms of memory (Diana et al., 2007; Eichenbaum et al., 2012; Eichenbaum et al., 2007; Yonelinas et al., 2022).

Sensory preconditioning is a form of second-order conditioning involving the integration of multiple associations formed through separate experiences (Brogden, 1939). In the preconditioning phase, subjects receive paired presentations of two neutral stimuli. Next, in the conditioning phase, subjects undergo first-order Pavlovian conditioning between one of the neutral stimuli and an unconditioned stimulus (US); this neutral stimulus hence becomes a conditioned stimulus (CS). Finally, in the test phase, the other neutral stimulus (the preconditioning stimulus, PCS) is presented to the subject. The PCS is expected to elicit the conditioned response, despite never having been directly paired with the US.

Sensory preconditioning may be supported by two cognitive mechanisms. The PCS may become directly associated with the US via “representation-mediated learning” (Holland, 1981). In this case, the CS evokes a representation of the PCS during conditioning, and an association is established between the PCS representation and the US. Alternatively, the PCS may elicit conditioned responding indirectly via an “associative chain” composed of the PCS-CS association formed during preconditioning and the CS-US association formed during conditioning (Rizley & Rescorla, 1972). Multiple factors seem to influence which mechanism supports a sensory preconditioning procedure (Holmes et al., 2022). However, we are not aware of any study that has systematically manipulated these factors and evaluated their influence.

Moreover, it is unclear whether different neural substrates are engaged by these two mechanisms. Both the PER (Holmes et al., 2013; Nicholson & Freeman Jr, 2000; Wong et al., 2019) and the POR (Taylor-Yeremeeva et al., 2021) are shown to be critically involved in sensory preconditioning. In these studies, however, the experimental procedures likely favored representation-mediated learning over associative chaining as the primary underlying mechanism. Therefore, whether the involvement of the PER and POR in sensory preconditioning extends to associative chaining is an open question.

In the current study, we utilized a sensory preconditioning procedure previously shown to rely on associative chaining at the time of PCS testing (Jones et al., 2012) and investigated the involvement of the PER and POR. Since we reasoned that PER and POR contributions to sensory preconditioning are the basis for their broader roles in latent learning (i.e. learning in the absence of explicit reward or punishment), we hypothesized that the two regions should be required for retrieving neutral associations for chaining as well. We therefore predicted reduced responding to the PCS when either the PER or the POR was temporarily unavailable during the PCS test. We employed the Designer-Receptor-Exclusively-Activated-by-Designer-Drugs (DREADDs) technique to create temporary suppression by systematic injection of clozapine-*N*-oxide prior to behavioral sessions (Smith et al., 2016; Urban & Roth, 2015).

## Materials and Methods

### Subjects

The subjects were 34 adult male Long-Evans rats (*Rattus norvegicus*) acquired from Charles River Laboratories (Wilmington, MA). Upon arrival at our facility, they were pair-housed in ventilated cages, placed on a reverse 12-hour light/dark cycle, and given *ad libitum* access to food and water. They were food restricted to 85-90% free-feed body weight starting from one week before surgeries until the end of the experiments. All subjects were about 3-4 months old at the time of surgeries and weighed between 300g to 400g. Subjects were handled by multiple lab personnel before surgery and during post-surgical recovery. Three subjects died during or shortly after long surgical procedures, one reacted badly to CNO injection procedures and was removed, and three subjects were removed due to equipment malfunction. This left 10 in the sham group, 11 in the PER group, and 9 in the POR group. All procedures were conducted according to NIH guidelines and protocols approved by the Brown University Institutional Animal Care and Use Committee.

### Surgery

The timeline for the procedures is shown in Figure 1 (panel A). Subjects were randomly assigned to the PER group, POR group, or Sham group before surgery. Pre-operative preparation included injections of glycopyrrolate, diazepam, and buprenorphine, and the subjects’ heads were shaved to expose the surgery site. Subjects were anesthetized with isoflurane and secured with ear bars in a stereotaxis apparatus. The bregma and lambda were arranged in the same horizontal plane (+/-0.1 mm) by adjusting the incisor bar. Craniotomies over the injection coordinates were made using a dental drill, and the dura was pierced prior to pipette insertion. For rats in the PER and POR group, bilateral injections of the DREADD virus AAV8-CaMKIIa-hM4d(Gi)-mCherry were made using a Hamilton syringe pump at 0.1 µl/min for 5 minutes per site (Table 1). The pipette tip remained at the injection coordinate for an additional 10 minutes before slowly being retracted. Rats in the Sham group received craniotomies only. Afterwards, gel foam soaked in sterile saline was placed over the cortex, and the incision was sutured. All subjects were given at least 2 weeks to recover from the surgery before behavioral procedures began.

**Figure 1.**
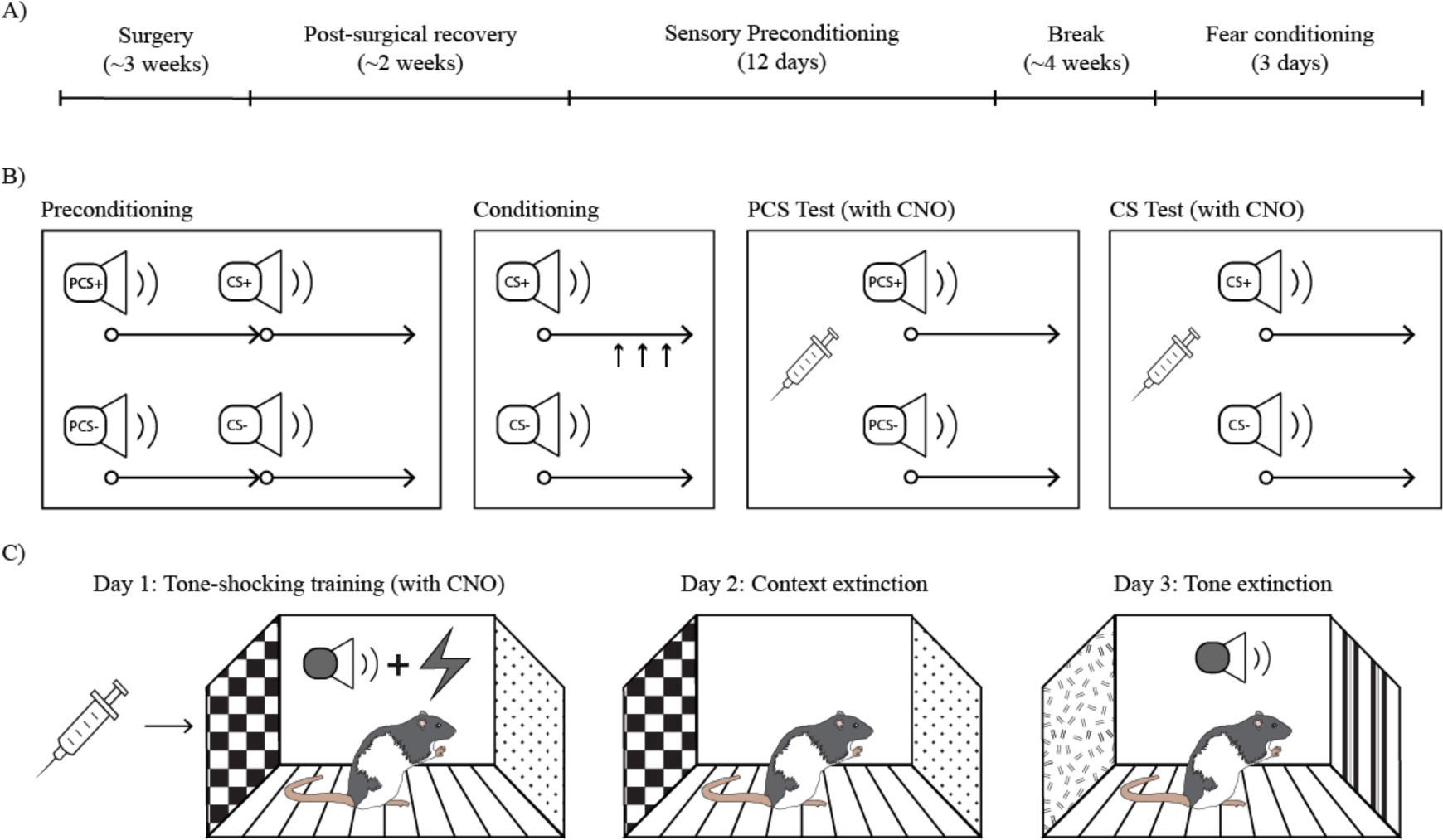
Schematics for experimental timeline and behavioral procedures. A) Overall timeline for the experiment, from the start of the surgeries until the end of behavioral testing. B) The Pavlovian sensory preconditioning procedure based on studies such as Jones et al. (2012) combined with CNO injections, indicated by the syringe symbol. Pellets were delivered only during the last 5 s of CS+ trials, indicated by upward arrows. C) A standard one-session fear conditioning procedure, followed with context extinction and tone extinction. CNO injections were only given on the first day.

**Table 1.** Stereotaxic coordinates for bilateral DREADD virus injections into PER or POR.

| Site | AP (mm) | ML (mm) | DV (mm) | Angle |
| --- | --- | --- | --- | --- |
| PER1 | 3.2 | +/- 5.0 | 6.3 | 13° |
| PER2 | 4.4 | +/- 5.0 | 6.2 | 13° |
| PER3 | 5.6 | +/- 5.0 | 5.9 | 13° |
| PER4 | 6.8 | +/- 5.0 | 5.5 | 13° |
| POR1 | -0.5* | +/- 5.0 | 4.5 | 16° |
| POR2 | 0.2* | +/- 5.0 | 2.75 | 16° |

In the anterior-posterior axis (AP), positive values indicate locations posterior to the reference point, and negative values indicate locations anterior to the reference. The bregma was used as reference for PER sites, whereas the lambda was the reference for POR sites (marked by \*). Dorsal-ventral (DV) values indicate distances to the skull surface.

### Apparatus

Behavioral procedures were performed in conditioning chambers (Med Associates) placed inside sound attenuating cabinets. The chambers were made with two aluminum side panels, three Plexiglas panels as front, back, and the ceiling. A food port located on one of the side panels was used to deliver pellets and detect the frequency and duration of nose pokes via infrared sensors. A speaker and a mechanical relay were located on the same side panel near the roof to produce the auditory stimuli used in the behavioral experiments. Four auditory stimuli (about 75 dB) were used for sensory preconditioning. The white noise and click (0.5 Hz) were used as PCS+ and PCS-, whereas a tone (2.5 kHz) and a siren (alternating between 1 kHz and 4 kHz) were used as CS+ and CS-. The stimulus arrangement was counter-balanced across subjects. The floor was made of stainless-steel rods connected to a shock generator and scrambler that were used to deliver foot shocks for fear conditioning. Video cameras mounted behind the back Plexiglas panel captured the fear conditioning sessions and saved the videos to a hard drive. Additional cues were added to the chamber for the fear conditioning experiment to differentiate the new contexts from the base context. These context cues included laminated sheets with distinctive visual patterns and artificial odors, such as vanilla and almond.

### Behavioral procedures

#### Sensory preconditioning

The general procedure was based upon prior studies from Schoenbaum and colleagues (Hart et al., 2020; Jones et al., 2012; Sadacca et al., 2018). Subjects first underwent a magazine training session, in which a total of 20 pellets were delivered via a variable-interval schedule. For this and all following sessions, the inter-trial intervals varied pseudo-randomly from three to six minutes. The procedure contained four phases (Figure 1B). The first was the preconditioning phase. Subjects were placed individually into conditioning chambers for two daily sessions of 12 preconditioning trials each. Each trial contained a serial presentation of one of two pairs of auditory stimuli. No reward was delivered during this phase. Next, in the conditioning phase, subjects received six sessions of first-order Pavlovian conditioning, each with 12 trials. During the CS+ trials, three food pellets total were delivered into the food port at 5, 7, 9 seconds after CS+ onset. No pellets were delivered during CS-. The third phase was the PCS testing phase. About 30 minutes prior to testing, all subjects received an intraperitoneal injection of clozapine *N*-oxide (CNO, 5mg per 1kg of body weight) and were returned to the home cage. The first 6 trials were CS+ and CS-reminder trials identical to trials in the conditioning phase. Next, subjects were presented with 12 PCS probe trials arranged in four 3-trial blocks in which the PCS+ and PCS-were presented without rewards. Finally, 48 hours after PCS testing, a 12-trial CS probe session was conducted about 30 minutes after intraperitoneal CNO injections. In each trial, subjects were presented with the CS+ or CS without reward delivery. All trial types in each session were counter balanced.

#### Fear conditioning

At least 30 days after the CS test session, a fear conditioning procedure was conducted to behaviorally verify the efficacy of DREADDs inhibition (Figure 1C). Though the same conditioning chambers were used, each subject was assigned a chamber different from the one where they underwent sensory preconditioning. Multiple visual, tactile, and olfactory cues were also introduced into the chambers to distinguish them from previous sensory preconditioning contexts. On day 1 (shock training with CNO), subjects received one session of signaled fear conditioning 30 minutes after CNO injections. Subjects were placed into context A.

After three minutes, three pure tones (10s, 2 kHz, 75dB) were delivered, each co-terminating with a foot shock (1s, 1.0 mA). The inter-stimulus-interval was one minute. Subjects remained in the chamber for an additional minute after the last tone-shock pairing before being returned to their home cages. On day 2 (context extinction without CNO), subjects were again placed into context A for 9 minutes without tone or shock delivery. On day 3 (tone extinction without CNO), subjects were put into context B. After 2 minutes, they received a continuous tone presentation lasting for 7 minutes. The specific configurations of context A and B were counter-balanced among subjects. All sessions were recorded with Ethovision 15 and saved to a computer hard drive for analysis.

### Behavioral and statistical analysis

#### Sensory preconditioning

Custom MED-PC scripts were written to control the stimuli and generate output containing the timestamps of all nose pokes into the food port. The output file was then analyzed by a custom-written Python script to compute the latency and total duration of nose pokes during periods of interest. For preconditioning, the nose poke time percentage differences were calculated between PCS+ and PCS-, and between CS+ and CS-. A positive difference score indicated a higher nose poke time percentage during the PCS+ versus PCS-, or the CS+ versus CS-. Separate one-way ANOVAs were conducted to detect whether any initial preference existed before conditioning started. For conditioning trials in the conditioning phase and CS reminder trials in PCS test, pellets were delivered during the second half (5 s) of the CS+, therefore conditioned responding was assessed by comparing the nose poke time percentages during the first half of the CS+ and CS-. For PCS and CS test trials, nose poke time percentages during the second half of the stimuli were compared. The difference scores were computed. First, a mixed-design ANOVA was conducted on difference scores from the conditioning sessions, with GROUP (Sham, PER, POR) as the between-subject factor and SESSION (1 to 6) as the within-subject factor. This test examined whether any group differences occurred during first-order Pavlovian conditioning, even though no CNO injection was made prior to the conditioning sessions. Second, a one-way ANOVA with GROUP as the between-subject factor was conducted on difference scores from PCS trials in the PCS test session under inactivation. Further, one-sample t tests were conducted to compare each group’s PCS difference score to 0. The Bonferroni-Holm procedure was applied to control for family-wise type I error rate (Abdi, 2010). These tests provided additional evidence on whether sensory preconditioning was successful in the Sham group, and whether suppression of the PER or POR led to deficits. The same analysis was conducted for CS test trials, also under DREADD suppression. These tests evaluated whether first-order conditioning was successful in the Sham group, and whether suppression of the PER or POR led to deficits. Finally, response latencies during PCS+ and CS+ trials were tested for group differences with a one-way ANOVA to determine whether attentional deficits contributed to group differences in nose poke time percentages.

#### Fear conditioning

The sessions were broken into 1-minute blocks, then a motion analysis software, custom-written in Python, was used to measure the total duration of freezing within each minute. Briefly, adjacent video frames were contrasted and processed to detect changes in pixel intensity that indicated movement from the subject. The subject was determined to be freezing when the period without movement exceeded a preset threshold. This software was validated with human observation in prior fear conditioning experiments (data not included). Expression of contextual fear was evaluated by analyzing freezing behavior on day 2 (context extinction). A mixed-design ANOVA was conducted with BLOCK as the within-subject factor and GROUP as the between-subject factor. Post hoc two-sided Dunnett’s t tests were used to compare each of the PER and POR group to the Sham group. Tone-elicited fear behavior on day 3 (tone extinction) was examined using the same tests.

### Histology

After the last fear conditioning session, the subjects were deeply anesthetized with isoflurane and received an overdose of Beuthanasia-D (100mg/kg, i.p.). They were then perfused with phosphate-buffered saline, followed by a 10% formalin solution. The brains were post-fixed for at least 48 hours in formalin and then were transferred to a 30% sucrose solution for at least another 48 hours. The brains were sectioned on a microtome into three series of 40μm sections. For each brain, one series was later stained with thionine, and another was stained with DAPI for fluorescence microscopy. Fluorescent images were taken in blue and red channels showing DAPI-stained neurons and neurons transfected by the DREADD virus, respectively. For histological analysis, each brain was scored based on expression in the rostral and caudal halves of the target area and on the degree of extra-target spread.

## Results

### Histology

Of the 11 rats in the PER group, viral transduction was observed bilaterally, and along the rostrocaudal extent of the target region in eight (Figure 2A). Thus, three were removed for insufficient transduction. For two of the eight cases, there was also moderate involvement in the ventral temporal association cortex and caudal hippocampus in both hemispheres, and these cases were also removed, leaving six in the PER group. Of the 9 rats in the POR group, viral transduction was observed bilaterally, and along the rostrocaudal extent of the region in seven (Figure 2B). For two cases, there was insufficient transduction, and those cases were removed. For three cases of the seven, there was slight involvement in the most caudal PER and/or caudal entorhinal cortex, and these cases were retained. In some cases, there was slight unilateral involvement of the temporal association cortex (TEv) dorsal to the POR, and these cases were also retained in the study.

**Figure 2.**
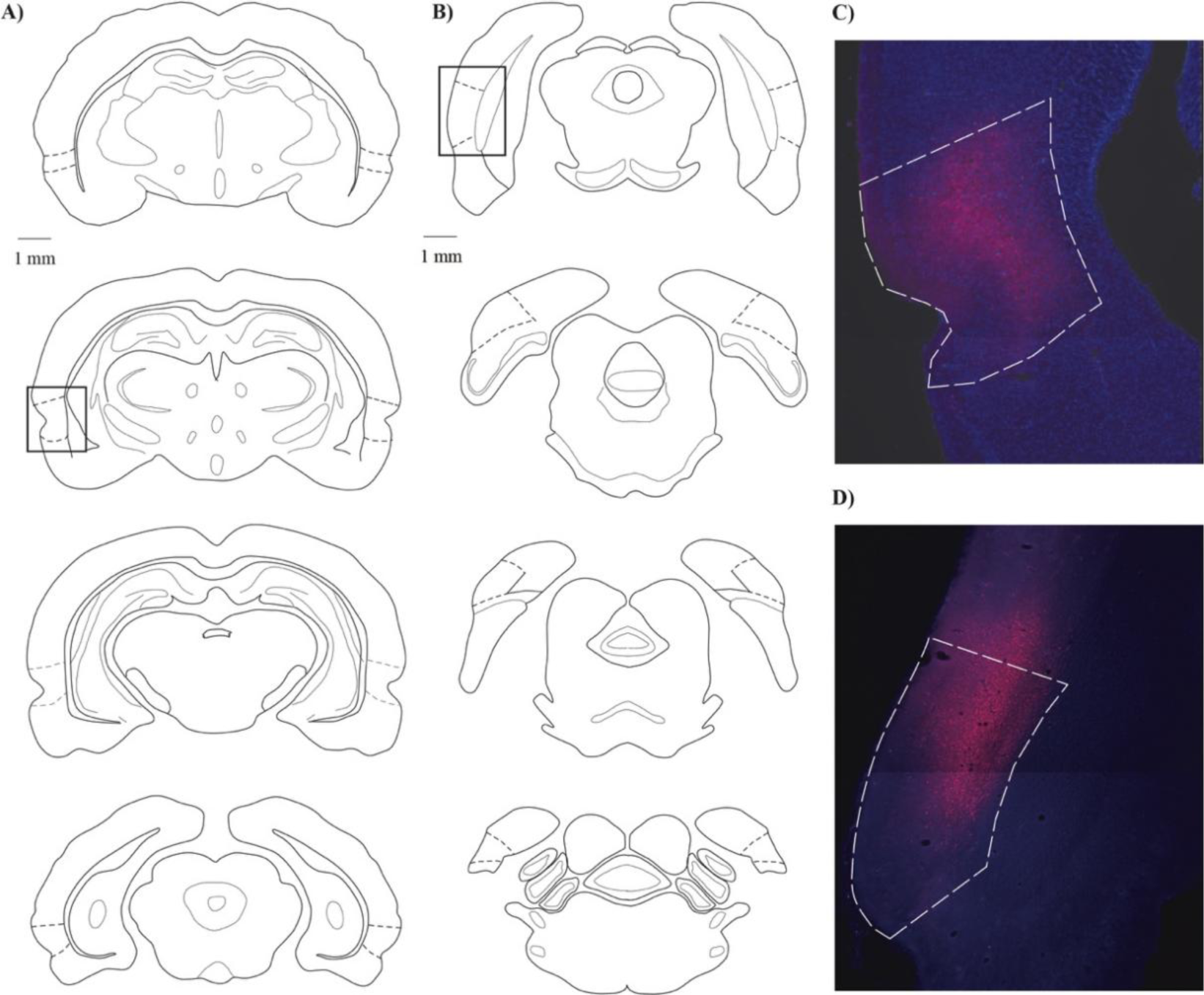
Schematics of PER and POR showing target locations along with representative labeling of viral transduction. A) Four levels of the PER with the target regions illustrated by dashed lines. Labeling was distributed across the rostrocaudal extent (see text for details). B) Four levels of the POR with the target regions illustrated by dashed line. Again, labeling was distributed across the rostrocaudal extent (see text for details). C) Example of labeling in PER. See inset in A for location. D). Example of labeling in POR. See inset in B for location. DAPI staining is captured in the blue channel, and the DREADD virus is captured in the red channel.

### Preconditioning and conditioning

The groups did not differ on nose poke time percentage differences during preconditioning sessions in their initial preference for PCS+ versus PCS-or CS+ versus CS-. A mixed-design ANOVA confirmed no significant DIFFERENCE*GROUP interaction (p = 0.792) and no main effects for DIFFERENCE (p = 0.625) or GROUP (p = 0.99). As expected, no impairment was found in any group for first-order conditioning (Figure 3). All groups significantly increased nose poke time during the first 5s of CS+, compared to CS-, and the changes were not significantly different among groups. A mixed-design ANOVA on nose poke time percentage differences for the 6 conditioning sessions showed a significant main effect for SESSION, F(5, 100) = 4.61, p = 0.001, but not for the SESSION*GROUP interaction (p = 0.247) or for GROUP (p = 0.849).

**Figure 3.**
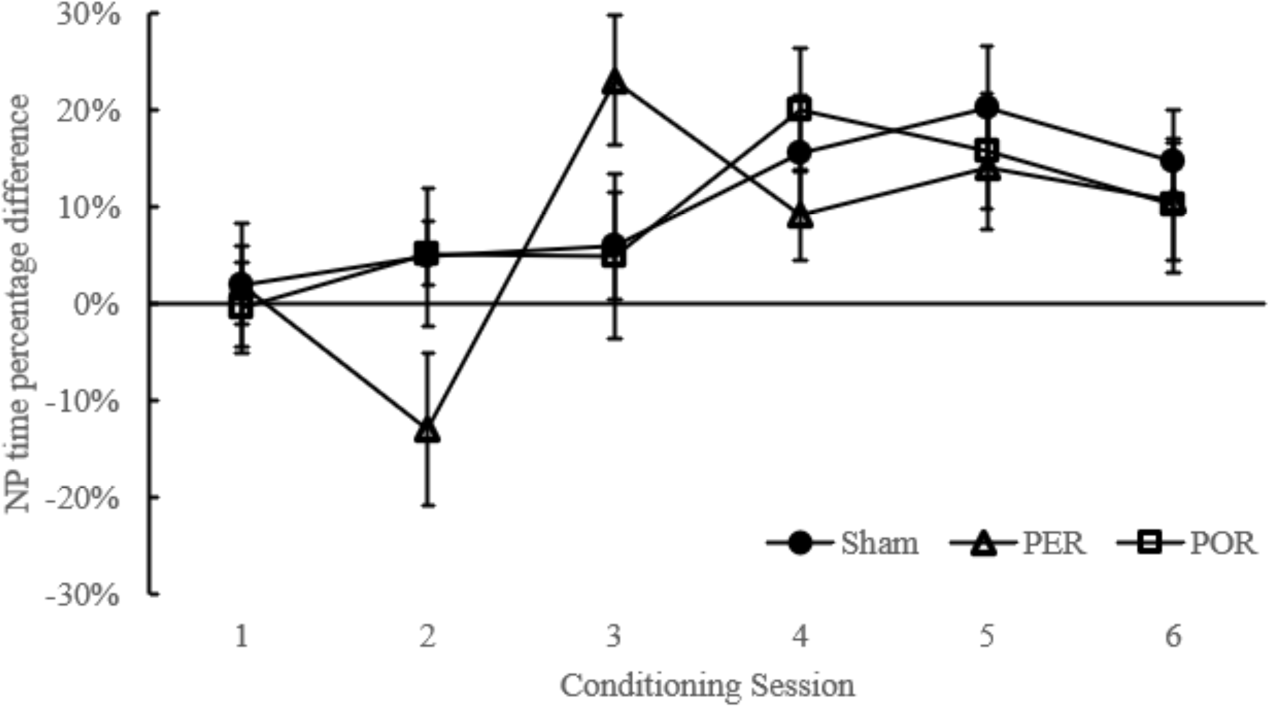
Nose poke time percentage difference during the conditioning phase. Data was only analyzed for the first half (5 s) of CS+ and CS-because pellets were delivered during the second half of CS+. Error bars indicate standard error.

### PCS test

The PCS test was conducted 30 minutes after the subjects received a CNO injection. The test started with 6 reminder trials that were identical to conditioning trials in that pellets were delivered during the last 5s of the CS+. The main measure for the sensory preconditioning effect was nose poke time the percentages during PCS+ and PCS-(Figure 4A). As predicted, the sham group showed a robust response sensory preconditioning effect, but the PER and POR groups did not (Figure 4). A one-way ANOVA was conducted to compare the nose poke time percentage difference between the last 5s of PCS+ versus PCS-. There were significant differences among groups, F(2, 20) = 3.82, p = 0.039 (Figure 4B). Planned comparisons revealed a marginally significant difference between the Sham group and the PER group, t(20) = 2.02, p = 0.057, and a significant difference between the Sham and the POR group, t(20) = 2.53, p = 0.020. Further, one-sample t tests showed that only the sham group had a significantly positive nose poke time percentage difference, t(9) = 3.71, p = 0.005. Neither the PER, t(5) = 0.97, p = 0.378, nor POR group t(6) = 0.001, p = 0.999, was significantly different from 0. Thus, after CNO injections, the sham group had significantly higher responding to PCS+ than PCS-during the last 5 s, whereas the PER and POR groups did not. This result provided evidence for an impairment in retrieving the PCS-CS association when either the PER or POR is under temporary DREADDs suppression.

**Figure 4.**
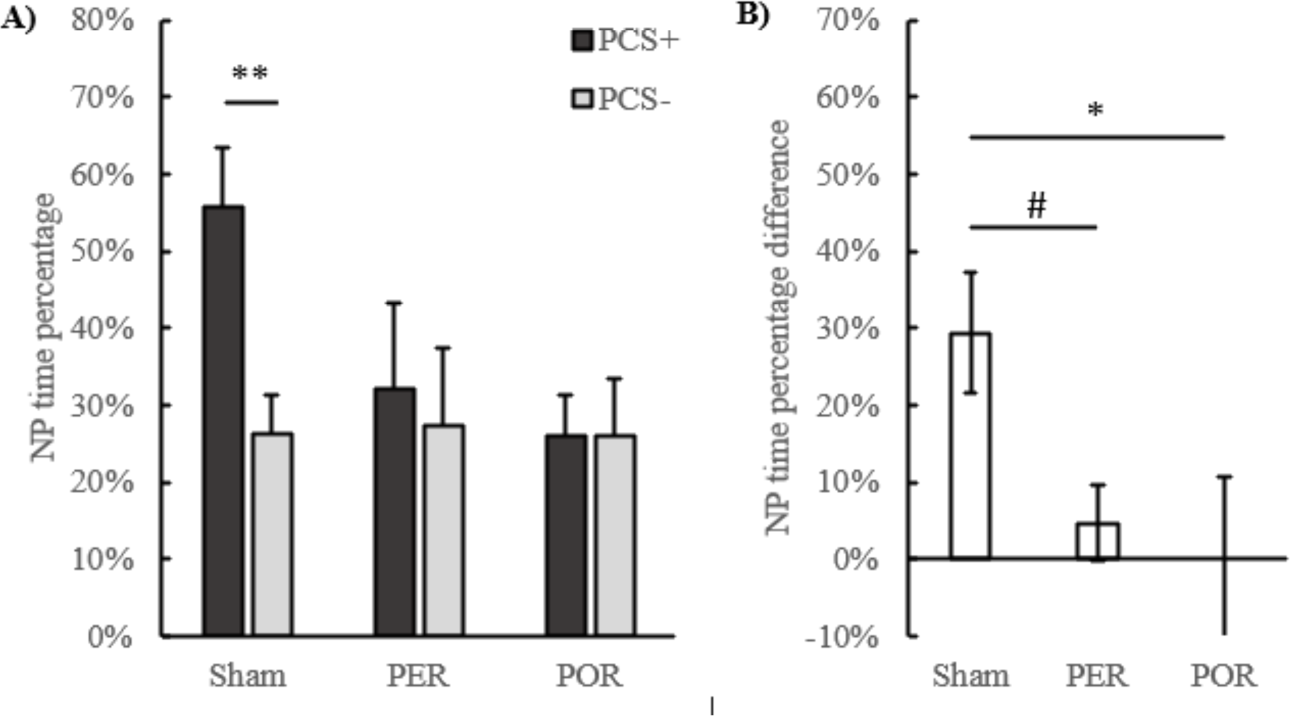
Nose poke time percentage during the last 5s of PCS trials in the PCS test phase with DREADD suppression. A) nose poke time percentages during PCS+ and PCS-. P-value symbols indicate significant differences between stimuli. B) nose poke time percentage difference between responding during PCS+ and PCS-. Positive values indicate higher responding during PCS+. P-value symbols indicate significant differences between groups. #: p < 0.10, *: p < 0.05, **: p < 0.01, ***: p < 0.001. Error bars indicate standard error.

### CS test

To assess first-order conditioning, a CS test session was conducted 48 hours after the PCS test, also following CNO injections. All groups showed a longer nose poke time percentage during the last 5s of the CS+, compared with the last 5s of the CS-(Figure 5A). The percentage time differences between CS+ and CS-were analyzed with one-way ANOVA (Figure 5B). The omnibus ANOVA was not significant, F(2, 20) = 2.39, p = 0.117. Planned comparisons revealed no significant difference between the Sham and PER groups, t(20) = 1.66, p = 0.112), but a marginally significant difference between the Sham and POR groups, t(20) = 2.01, p = 0.064. One-sample t tests showed that all groups had significantly positive difference scores during the CS test: sham, t(9) = 7.96, p < 0.001; PER, t(5) = 3.40, p = 0.019; POR, t(6) = 6.79, p < 0.001. Overall, all groups demonstrated significantly longer nose poke time percentage during the CS+, compared with the CS-, indicating successful first-order conditioning.

**Figure 5.**
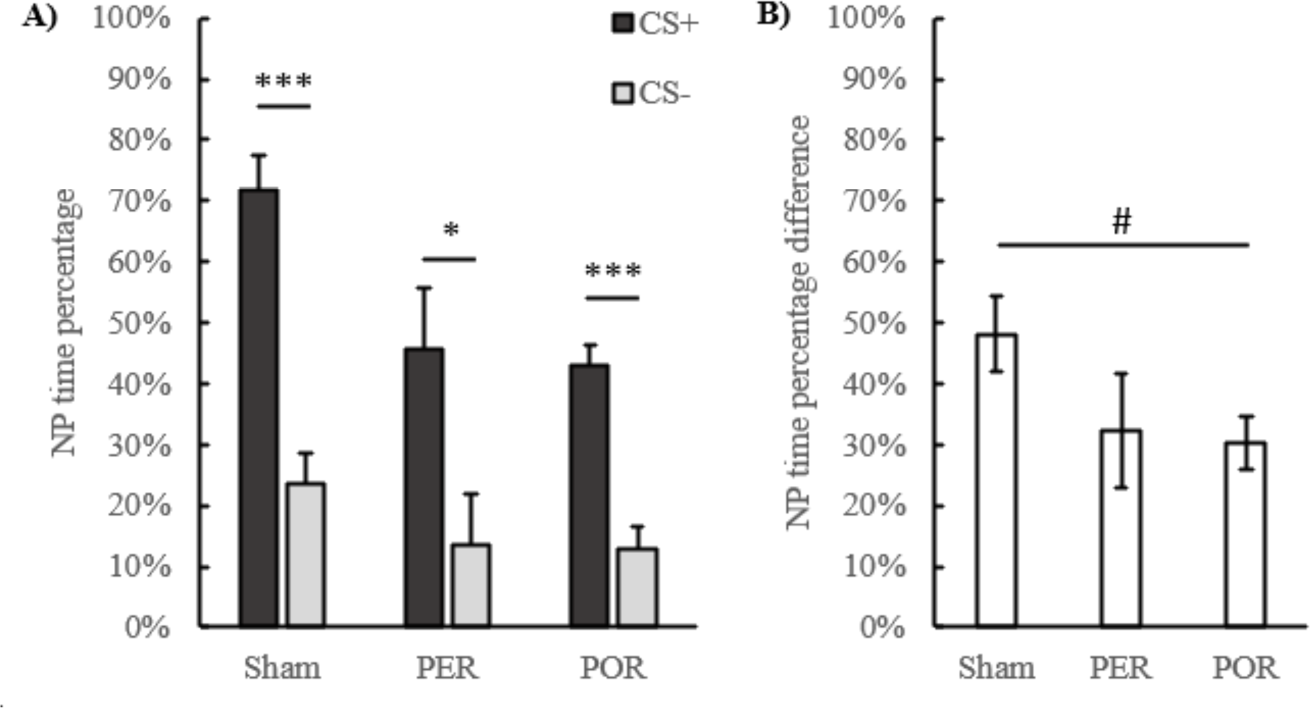
Nose poke time percentages during the last 5s of CS trials in the CS test with DREADD suppression. A) nose poke time percentages during CS+ and CS-. Asterisks indicate significant one-sample t-tests on the difference between CS+ and CS-. B) nose poke time percentage differences between CS+and CS-. Error bars indicate standard error.

### PCS and CS response latency

To confirm that DREADD suppression did not lead to attentional deficits in the PER or the POR group, nose poke latencies after stimulus onset during the PCS+ and CS+ test trials were analyzed separately with one-way ANOVA (Figure 6). The omnibus ANOVA was not significant for CS+ trials (p = 0.308) or PCS+ trials (p = 0. 379), and none of the planned comparisons between the Sham and virus groups was significant. Therefore, deficits observed in nose poke time percentages cannot be explained by delayed reaction in response to stimulus onset.

**Figure 6.**
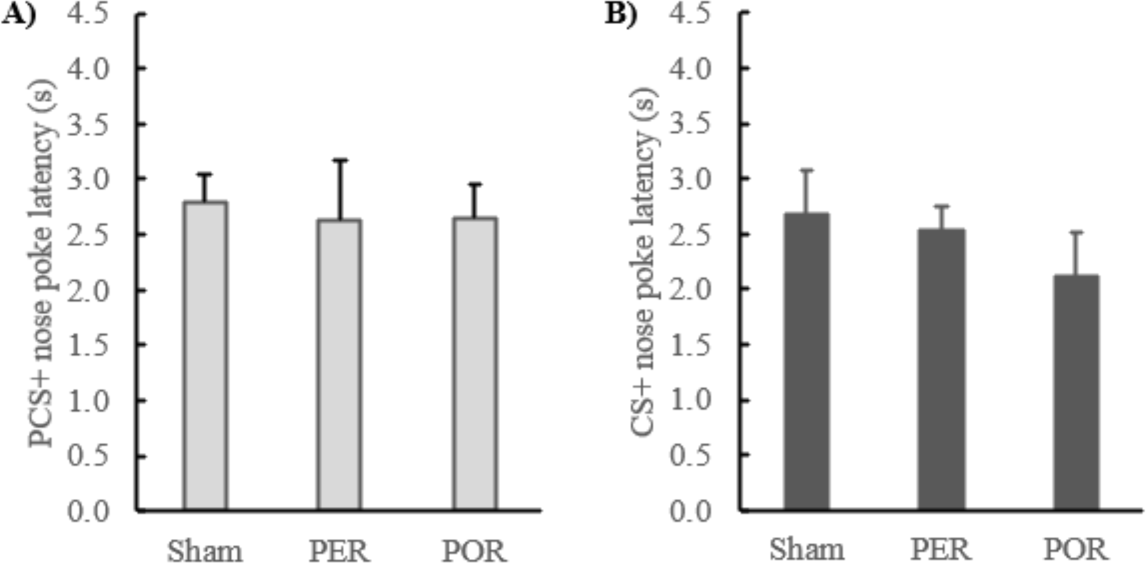
Latencies of nose poking in the test sessions under suppression. A) nose poke latencies since the onset of PCS+ during test trials. B) nose poke latencies since the onset of CS+ during test trials. Error bars indicate standard error.

### Fear conditioning

In a fear conditioning paradigm, subjects received an injection of CNO (i.p.) on day 1, followed by tone-shock conditioning in context A. On day 2, they received extinction in context A without the tone. And on day 3, they underwent tone extinction in context B. No CNO injections were made on day 2 or 3. For freezing time on day 2 (Figure 7A), a mixed-design ANOVA did not reveal a significant MINUTE*GROUP interaction, F(16, 160) = 0.968, p = 0.494. However, there was a significant main effect of GROUP, F(2, 20) = 6.87, p = 0.005, as well as a main effect of MINUTE, F(4.9, 160) = 2.47, p = 0.039. Further simple contrast tests revealed a significant difference between the Sham and the PER group, p = 0.002, but not between the Sham and the POR group, p = 0.79. Therefore, when re-exposed to the shock context, the PER group displayed a significantly lower amount of freezing when compared to the sham group, indicative of impaired contextual fear memory. No reduction in freezing was observed in the POR group, indicating normal contextual fear memory. Since DREADDs suppression was induced only during tone-shock training (day 1) but not on day 2, this suggests that the acquisition of contextual fear was impaired in the PER group, but not in the POR group, when compared to the Sham group.

**Figure 7.**
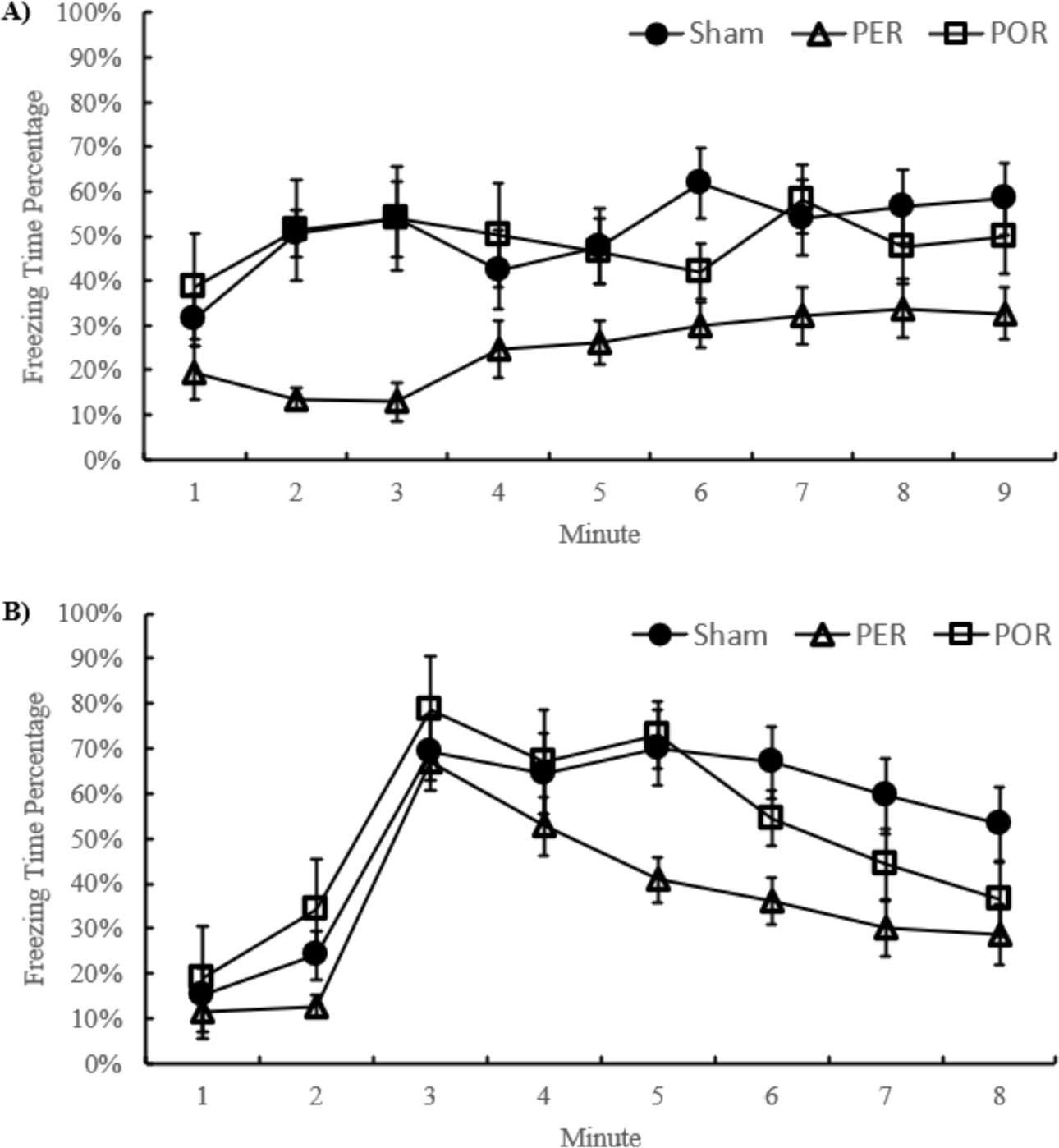
Freezing time percentage in each minute during context extinction (day 2, panel A) and tone extinction (day 3, panel B). The tone was presented from the start of the third minute during tone extinction. Error bars indicate standard error.

Freezing to the tone on day 3 was also analyzed by a mixed-design ANOVA to detect the presence of tone-elicited fear (Figure 7B). A significant main effect of MINUTE was revealed, F(3.74, 74.81) = 11.76, p < 0.001, but no significant MINUTE *GROUP interaction, F(7.48, 74.81) = 1.34, p = 0.203, or main effect of GROUP, F(2, 20) = 1.77, p = 0.196. Simple contrast tests revealed a marginally significant difference between the Sham and the PER group (p = 0.82). Further tests revealed a significant linear trend for MINUTE, F(1, 20) = 36.60, p < 0.001. The results indicated that all three groups gradually displayed less freezing to the tone, with no significant group difference. Therefore, fear conditioning to the tone was not impaired by PER or POR suppression during conditioning.

## Discussion

Chemogenetically suppressing bilateral PER or POR during the PCS test impaired sensory preconditioning by disrupting the retrieval of PCS-CS associations. First-order conditioning remained intact in all groups under the influence of CNO. The overall results suggest that, although the PER and POR are not necessary for conditioned responding to directly conditioned auditory stimuli used in this experiment, both are required for retrieving neutral stimulus-stimulus (S-S) associations when re-exposed to the PCS. These findings appear partially inconsistent with existing studies using temporary PER or POR suppression. However, we argue that this is because the sensory preconditioning effect in our study relied on a mechanism different from procedures in previous studies. Whereas existing studies collectively showed that the PER and POR are necessary for sensory preconditioning when it primarily depends on representation-mediated learning, our findings indicate they are crucial for associative chaining as well.

Our sensory preconditioning procedure was slightly modified from a previous study in which sensory preconditioning deficits occurred when temporarily suppressing the orbitofrontal cortex during the PCS test (Jones et al., 2012). This pattern of deficit typically indicates that responding to the preconditioning cue is likely supported by associative chaining (Holmes et al., 2022; Jones et al., 2012). In contrast, previous studies of the PER or POR employed procedures that likely relied on representation-mediated learning (Holland, 1981). A recent study found reduced responding to the PCS if chemogenetic POR suppression took place during preconditioning or conditioning, but not if it occurred during PCS testing (Taylor-Yeremeeva et al., 2021). In Wong et al. (2019), reduced sensory preconditioning was observed when a sodium channel blocker was injected into the PER before conditioning or before the test (albeit of a smaller degree). The authors concluded that PER-dependent memory integration in their procedure occurred primarily via mediated learning during conditioning, though it wasn’t ruled out that some associative chaining also took place at the time of testing. Collectively, these studies and ours suggest the PER and POR are necessary for retrieving neutral S-S associations, regardless of whether these associations are used for mediated learning or associative chaining. Moreover, both mechanisms were shown to be necessary for encoding associations between neutral stimuli during preconditioning (Holmes et al., 2013; Taylor-Yeremeeva et al., 2021). Overall, the evidence supports the notion that the critical contribution of the PER and POR to sensory preconditioning is the encoding and retrieving of the associations formed when the stimuli involved are neutral.

The ability to associate neutral stimuli in the absence of explicit reward or punishment fits well with the roles of the PER and POR in automatically representing the spatial context and using contextual cues to guide spontaneous or learned behavior (Peng & Burwell, 2021). At the most basic level, the context exists as a loose collection of environmental elements that contain multi-sensory information without meaningful connections. But with the contribution from MTL structures such as the PER and POR, associations formed among the elements become the basis for a cohesive representation of the context. For example, neurons in the POR increase firing rates when the rat encounters a particular object at a particular location, but not when encountering the same object at other locations or other objects at the same location (Furtak et al., 2007). This “object-location” conjunctive coding likely emerges as object information from the PER is combined with spatial information from POR via robust direct interconnections (Agster & Burwell, 2013; Heimer-McGinn et al., 2017). The PER and POR are necessary in a variation of sensory preconditioning in which the preconditioning phase requires establishing an association between a neutral cue and the context (Iordanova et al., 2009; Taylor-Yeremeeva et al., 2021).

In typical context learning paradigms, the context is usually composed of static stimuli that remain constant throughout a session. In contrast, associations formed and retrieved in sensory preconditioning are among transient stimuli with clear onsets and offsets. Therefore, the critical involvement of the PER and POR in sensory preconditioning suggests their contributions extend beyond the well-known ability for context learning; they may also be capable of representing transient cues inside the context. In other words, they are not only able to represent stimuli existing in the three-dimensional *space*, but also stimuli in the dimension of *time*. This is consistent with findings about events and sequences representations in other parts of the MTL, such as the entorhinal cortex (Robinson et al., 2017) and the hippocampus (Allen et al., 2016; MacDonald et al., 2013; Pastalkova et al., 2008). The PER is necessary for bridging the temporal gap between segments of a discontinuous auditory stimulus (Bang & Brown, 2009; Kholodar-Smith, Allen, & Brown, 2008) and between a CS and US in trace fear conditioning (Kholodar-Smith, Boguszewski, & Brown, 2008). It is likely supported by persistent firing PER neurons that can maintain activation for seconds to several minutes (Navaroli et al., 2012). The PER is also important for incidental learning of odor sequences (Allen et al., 2020; Jayachandran et al., 2019). The POR also plays a role in processing localized temporal events with a significant spatial component, in which case it may cooperate with the PER cortex via their functional connectivity (Heimer-McGinn et al., 2017).

Combining the evidence that both areas are necessary for incidental learning of event associations together with their well-established roles in representing context, we propose that the rodent PER and POR may collectively construct and maintain neural representations of stimulus sequences in contexts that serve as rudimentary forms of event representation. These representations contain salient features of an event, such as the occurrence of transient cues, their associations and temporal relationships, and the context. In other words, these “event elements” in the PER and POR constitutes the “gist”, but not the full perceptual aspects of an event (Moscovitch et al., 2016; Robin & Moscovitch, 2017). These representations may be available to the hippocampus as the “backbone” for episodic memory (Agster & Burwell, 2013; Furtak et al., 2007). Alternatively, they may also be used by areas outside the MTL for hippocampal-independent behaviors, such as expressing remote fear memory (Burwell et al., 2004), or when hippocampal functioning is disrupted (Fanselow, 2010; Wiltgen et al., 2006). This does not detract from the key role of the hippocampus in episodic memory (Eichenbaum, 2017; Eichenbaum et al., 2012; Moscovitch et al., 2016; Robin & Moscovitch, 2017). Instead, it fits into the idea that two types of episode memory representations dynamically co-exist in the brain, with one being “schematic” or “gist-like” that mostly resides in the neocortex, and the other one being detail-rich and hippocampal-dependent (Moscovitch et al., 2016; Robin & Moscovitch, 2017). The PER and POR have reciprocal connections with the hippocampus and various prefrontal areas (Agster & Burwell, 2013; Furtak et al., 2007; Hwang et al., 2018), therefore they potentially serve as a region where the two types of episodic representations overlap or interact. Other cortical areas such as the orbitofrontal cortex and the retrosplenial cortex may be involved as well (Hart et al., 2020; Jones et al., 2012; Robinson et al., 2014; Todd et al., 2019).

Results from the contextual fear conditioning with DREADD inactivation during training were partially inconsistent with existing literature. Prior studies found that normal contextual fear learning required both the PER and POR (Bucci et al., 2000; Bucci et al., 2002). Although in the current study the PER group displayed less freezing during the context extinction test as expected, the POR group had similar levels of context-elicited freezing compared to the Sham group. We argue this discrepancy regarding effects of POR inactivation was because the fear conditioning contexts were not sufficiently distinctive from the sensory preconditioning context, even though we did introduce various visual and olfactory cues to the chambers for fear conditioning. Consequently, unlike typical fear conditioning in which the context is novel to the subjects, our subjects were already partially familiar with context where shocks occurred. This may suggest that although contextual representations initially depend on both the PER and POR, as the context becomes increasingly familiar, the PER alone gradually becomes more efficient at representing the context thanks to visuospatial input from the POR and other areas, until the PER alone may be sufficient for new learning, such as contextual fear conditioning.

In conclusion, we utilized the DREADD technique to show that both the PER and POR are necessary for retrieving neutral S-S associations in a sensory preconditioning procedure that likely depends on the chaining of associations at PCS testing. Combined with their well-established roles in context processing, these findings suggest the PER and POR collectively support a rudimentary, gist-like representation of event elements that could form the basis for episodic-like memory in rodents. Although latent learning paradigms such as sensory preconditioning do not capture the full spectrum of episodic-like memory in animals (Clayton et al., 2003), they nevertheless provide us with a powerful tool to investigate the neural mechanisms underlying different aspects of episodic-like memory in animal models.

## Notes

### Competing Interest Statement

The authors have declared no competing interest.

